# Context-bridged associative learning: linking neutral tone engram to fear through shared context

**DOI:** 10.64898/2026.02.26.708189

**Authors:** Olga I. Ivashkina, Ksenia A. Toropova, Konstantin Anokhin

## Abstract

In associative fear learning, weak or temporally constrained training may fail to link a conditioned stimulus (CS) with an aversive unconditioned stimulus (US), particularly when the contextual representation is impaired (the immediate shock deficit). Here, we systematically tested behavioral conditions that enable linking of an initially neutral auditory memory trace to an aversive episode. Male C57BL/6 mice were studied in four experiments manipulating (i) preexposure to a CS tone, (ii) the duration of context exploration before immediate footshock, and (iii) whether CS memory was tested in a novel or a familiar-like context. A 5 s tone followed immediately by footshock did not induce reliable fear to either the CS or the training context. CS preexposure three days before conditioning did not facilitate CS aversive memory when animals were tested in a completely novel context. However, robust facilitation emerged when the CS memory was tested in a context similar to the preexposure/conditioning one, indicating strong contextual gating of CS retrieval. Extending context exploration before shock enabled CS fear learning, but reduced (and even reversed) the effect of CS preexposure, consistent with latent inhibition. Together, these results delineate behavioral constraints for linking an initially neutral cue memory to an aversive event and highlight contextual control over the coupling and expression of cue memory traces.

## 1. Introduction

Fear conditioning provides a tractable model to study how neutral cues and contexts become associated with aversive outcomes and how such memories generalize across environments. Depending on temporal relation and training parameters, Pavlovian conditioning can produce strong and persistent conditioned freezing to both discrete conditioned stimuli (CSs) and to the conditioning context itself [1-5]. In rodents, hippocampal representations of context interact with amygdala and prefrontal circuits to control retrieval, generalization, and renewal of fear [6-11]. A key challenge is to understand how an initially non-aversive cue representation becomes integrated with aversive stimulus, and under which conditions this integration is context-specific. Preexposure can exert opposite effects depending on what is preexposed and for how long: lengthy preexposure to a discrete CS can reduce subsequent conditioning (latent inhibition) [12-13], whereas short preexposure to the context can facilitate subsequent contextual fear learning under immediate-shock conditions (the context preexposure facilitation effect - CPFE) [14-16]. In standard immediate-shock (IS) procedures, little fear is acquired because the animal has insufficient time to encode a conjunctive (configural) representation of the context before the aversive event [17]. The CPFE refers to the robust enhancement of contextual fear conditioning when animals are familiarized with the training context and only later receive an IS there [14-16]. Context preexposure supports hippocampus-dependent formation (and later retrieval) of a configural context representation, which can then be linked to shock during IS training and expressed as robust freezing at test [7, 18, 19]. CPFE is therefore widely used as an assay of hippocampus-dependent context representation, pattern completion, and the temporal separation of context encoding from shock association [6, 9]. Under appropriate conditions, CPFE also provides a window into factors that modulate context encoding and its later integration with aversive outcomes [20-21].

Here we focused on whether a neutral tone representation, formed during preexposure, can be linked to aversive stimulus when an IS follows the tone and when subsequent testing occurs in contexts that vary in similarity to the original preexposure environment. IS designs are informative because they probe how rapidly contextual or cue information is acquired and how cue and context components compete during learning and retrieval [4, 14-17].

At the circuit level, these questions can be framed in terms of how conjunctive representations and pattern completion in hippocampal networks support context-dependent retrieval of compound experiences [6, 9, 22]. Context similarity at test may therefore gate which components of a prior episode are reactivated, biasing expression toward contextual fear or toward cue-driven responding. Recent engram frameworks further suggest that cross-event linking can arise through biased allocation to overlapping neuronal ensembles driven by transient changes in intrinsic excitability and epigenetic priming [23-26]. Mechanisms that preserve ensemble stability and constrain overgeneralization are increasingly recognized as key determinants of memory precision across hippocampal and cortical circuits, providing an additional mechanistic backdrop for context-dependent fear expression [27, 28].

In this study, we used conditioning protocols that varied the duration of context exploration and the availability of tone preexposure to determine when the cue memory trace is linked to an IS and how context similarity at test modulates fear expression.

## 2. Results

### 2.1. A 5 s Tone with Immediate Shock Does Not Produce Reliable Cue or Context Fear

First, we tested whether an extremely brief tone–shock pairing (5 s tone followed immediately by FS) is sufficient to form cued associative memory, analogous to classical cue fear conditioning, despite minimal time for contextual processing.

No significant differences between groups were detected during the tone test in Context B or during the context A test (Figure 1). Freezing levels were very low both in Context B and during tone presentation (means 3.03% and 4.40%, respectively), indicating that this procedure did not establish detectable associative memory for the tone. During the Context A test, the average freezing was higher (18.66%). However, as the active control group (A + Tone) never received FS, this freezing cannot be attributed to conditioned fear of the context and likely reflects non-associative immobility rather than a learned context response. Thus, consistent with IS-related literature on contextual learning, a 5 s exposure paired with immediate shock was insufficient not only for contextual fear but also for forming a robust associative memory for a discrete auditory cue under these conditions.

**Figure 1.**
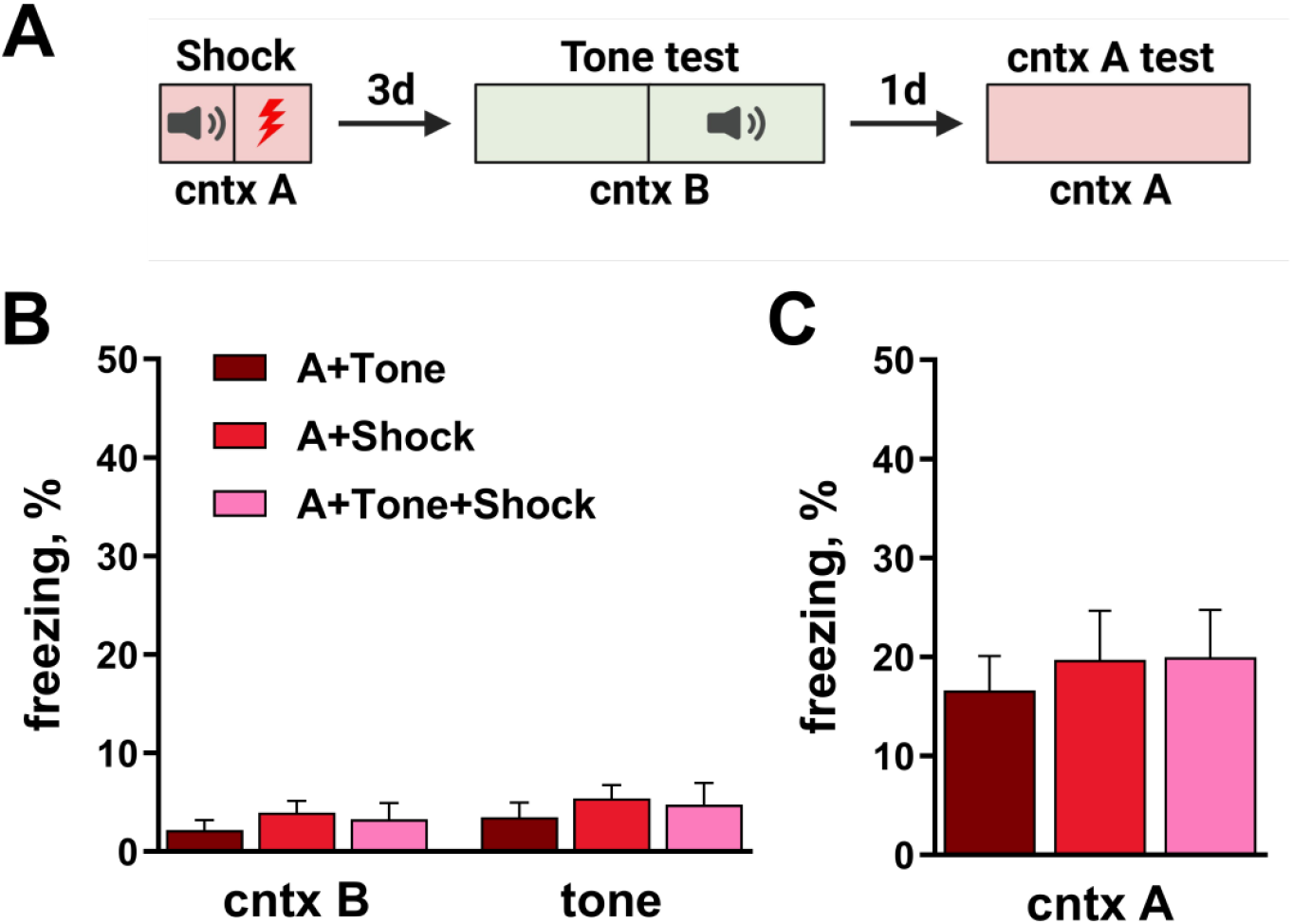
A 5 s tone paired with immediate shock does not induce fear to the tone. (A) Experimental design. (B) Freezing in Context B during baseline exploration and during the tone presentation. (C) Freezing during the Context A test.

### 2.2. Tone Preexposure Does Not Facilitate Tone Fear When Tested in a Novel Context

We further asked whether preexposure to the tone in Context A (three 20 s presentations during a 5 min session) would facilitate subsequent conditioning when the tone was later presented for 5 s and paired immediately with FS.

During tone preexposure, freezing remained low across all groups (means 4.19% in Context A and 6.63% during tone presentations), with no significant group differences (Figure 2B), indicating that tone exposure did not elicit nonspecific fear.

**Figure 2.**
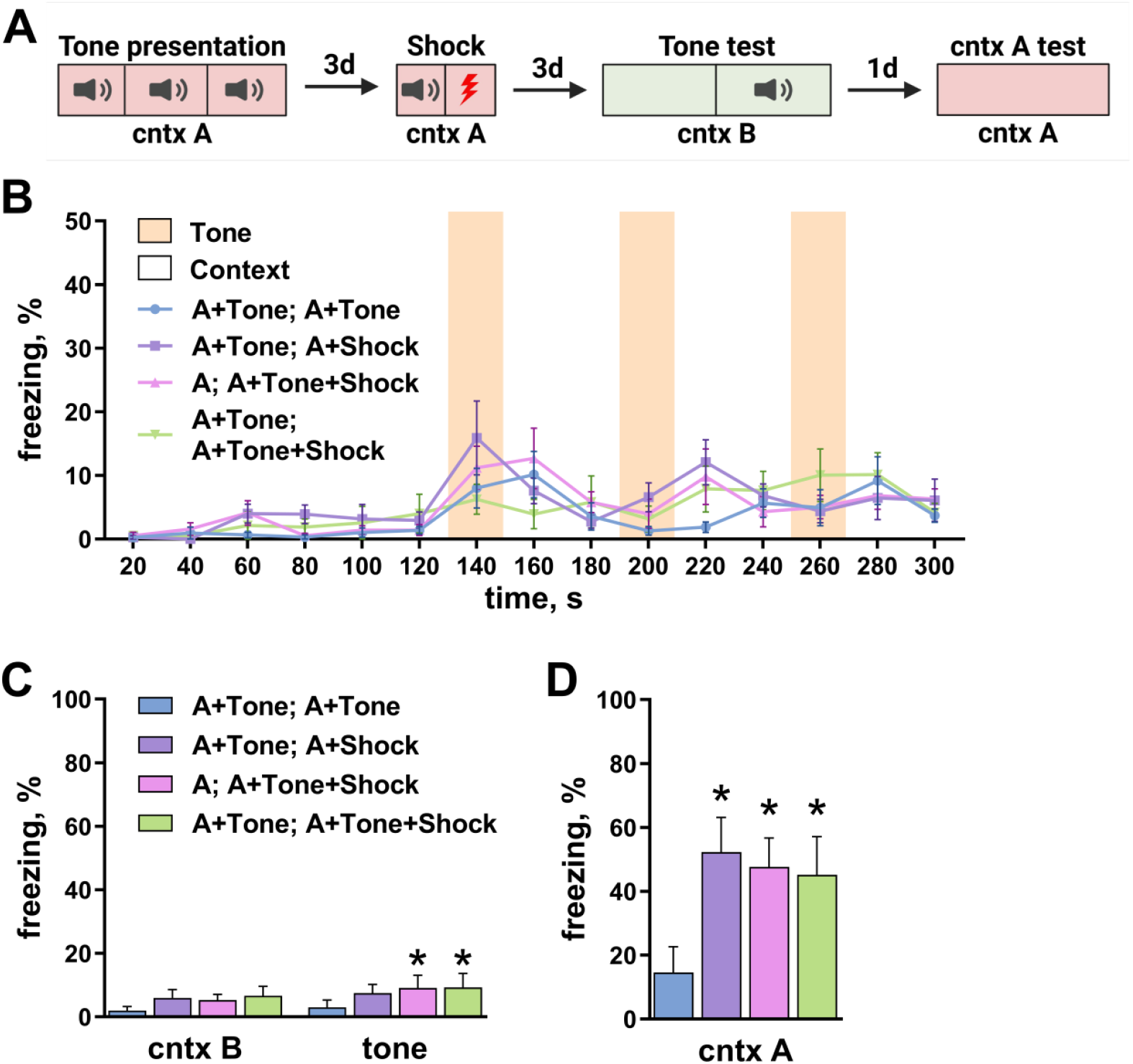
Preexposure to a tone does not facilitate subsequent tone fear when retrieval is tested in a novel context. (A) Experimental design. (B) Freezing in Context A during the tone preexposure session and during tone presentations; * p < 0.0003 compared to “A+Tone; A+Tone” group, Tukey’s post hoc test. (C) Freezing in Context B during baseline exploration and during the tone presentation (tone test). (D) Freezing during the Context A test; * - p < 0.0003 compared to “A+Tone; A+Tone” group, Dunnett’s T3 post hoc test.

During the tone test in Context B, freezing remained low across groups. A significant main effect of group was detected (F(3,34) = 4.984, p = 0.0057; test episode: F(1,34) = 9.661, p = 0.0038; Figure 2C), driven by higher baseline freezing in Context B in the A + Tone; A + Shock group compared with the A + Tone; A + Tone + Shock group (Tukey’s post hoc test, p < 0.05). Importantly, freezing during tone presentation did not show a selective increase in the conditioned group, and mean values were low overall (5.03% baseline in Context B and 7.34% during tone).

By contrast, during the context A test, all groups that had received shock displayed significantly higher freezing than the active control (A + Tone; A + Tone) (F(3,34) = 14.19, p < 0.0001; pairwise comparisons vs. the active control: p < 0.0003; Figure 2D). Importantly, presenting the tone during the preexposure session did not impair context conditioning: freezing did not differ between the shock groups (A + Tone; A + Shock; A; A + Tone + Shock; A + Tone; A + Tone + Shock).

Therefore, under immediate-shock conditioning and when tested in a novel context, tone preexposure did not facilitate the formation or expression of tone associative fear, whereas contextual fear was readily observed.

### 2.3. Tone Preexposure Facilitates Tone Fear in a Context-Dependent Manner

The negative result for cue fear in Experiment 2.2 could reflect either a failure to form tone fear or an inability to retrieve/express it in a completely novel test context. Therefore, in this experiment, mice were tested in Context A′, which shared key features with the conditioning context.

During the tone test in Context A′, all groups that received FS showed significantly higher freezing than the active control group, both during baseline exploration of A′ and during tone presentation (group: F(3,36) = 37.86, p < 0.0001; repeated measures: F(1,36) = 68.31, p < 0.0001; interaction: F(3,36) = 27.30, p < 0.0001). Post hoc comparisons confirmed higher freezing relative to the active control in both periods (baseline: p < 0.0017; tone: p < 0.0007; Figure 3B). Baseline freezing in A′ remained relatively modest (mean 17.23% across the groups with FS), consistent with partial similarity between A′ and the FS context A.

**Figure 3.**
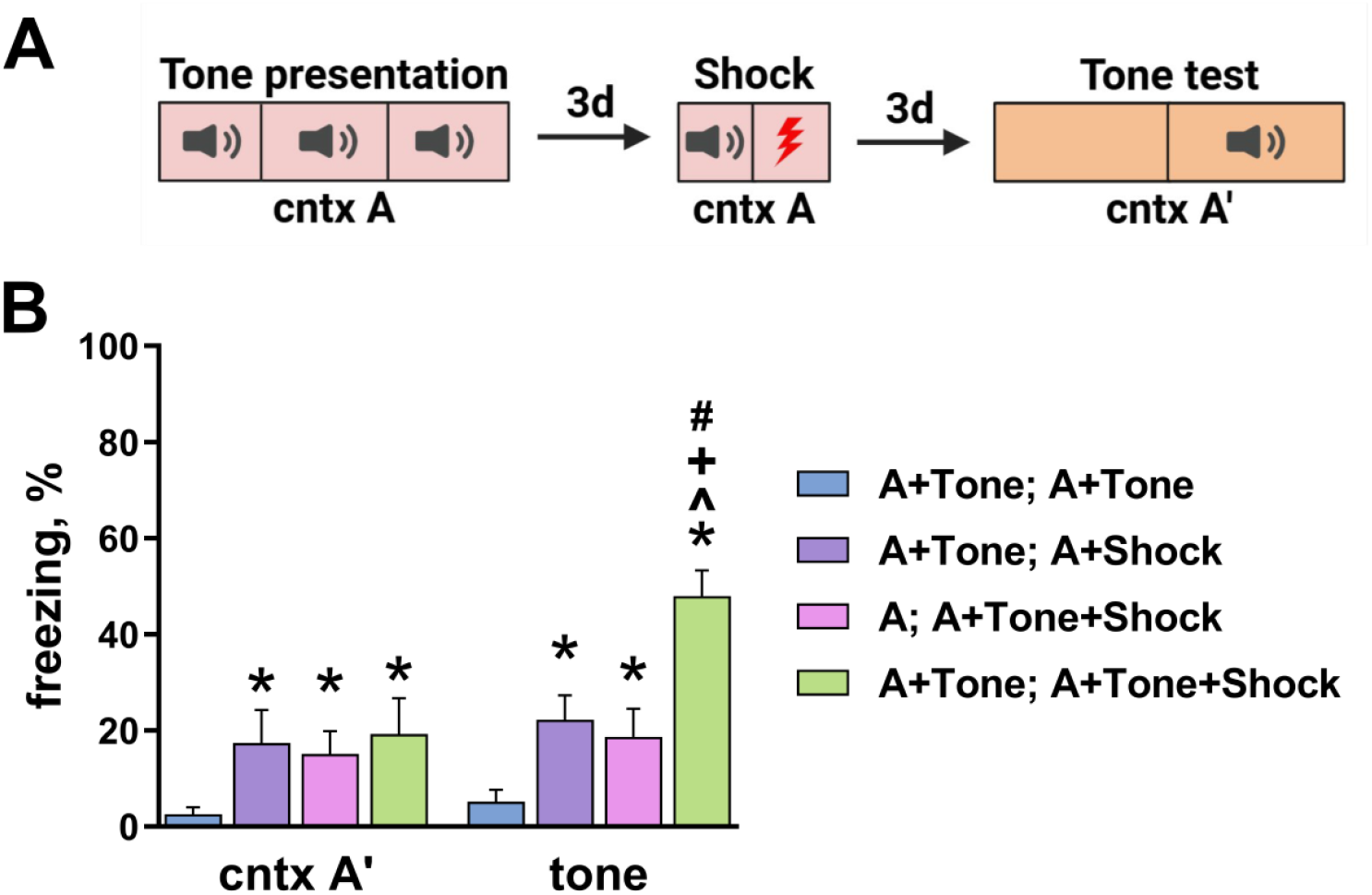
Tone preexposure facilitates tone fear when retrieval is done in a context similar to the conditioning context. (A) Experimental design. (B) Freezing in Context A′ during baseline exploration and during the tone presentation; * p < 0.0003 compared to “A+Tone; A+Tone”; + - p < 0.0001 compared to “A+Tone; A+Shock”; ^ - p < 0.0001 compared to “A; A+Tone+Shock”; # - p < 0.0001 compared to freezing in Context A’ within the same group, Tukey’s post hoc test.

Importantly, during tone presentation, the preexposed-and-conditioned group (A + Tone; A + Tone + Shock) showed the highest freezing: animals froze significantly more than in the other FS groups (p < 0.0001) and also significantly more than during baseline exploration of A′ (p < 0.0001). Thus, tone preexposure facilitated the formation and/or expression of associative fear to the tone, but this effect was evident only when retrieval was probed in a context similar to the one in which the tone was experienced and paired with FS. This indicates strong context dependence of cue fear under these training conditions.

### 2.4. Longer Context Exploration Enables Tone Fear but Reveals Latent Inhibition after Tone Preexposure

Finally, we tested whether extending the time available for contextual processing during conditioning (200 s total in Context A, with 150 s exploration before tone and shock) would reduce context dependence and enable tone fear without requiring tone preexposure.

During conditioning, freezing was low during the initial exploration period and during tone presentation, with no group differences in this pre-shock interval (two-way ANOVA: group: F(3,113) = 6.705, p = 0.0003; training episode: F(10,1130) = 171.0, p < 0.0001; interaction: F(30,1130) = 14.17, p < 0.0001; pairwise comparisons between groups in the 0–165 s interval: p > 0.1520; Figure 4B). After FS, freezing increased robustly and similarly in all shock groups compared to the active control (comparisons vs. A + Tone; A + Tone in the 170–200 s interval: p < 0.0001).

**Figure 4.**
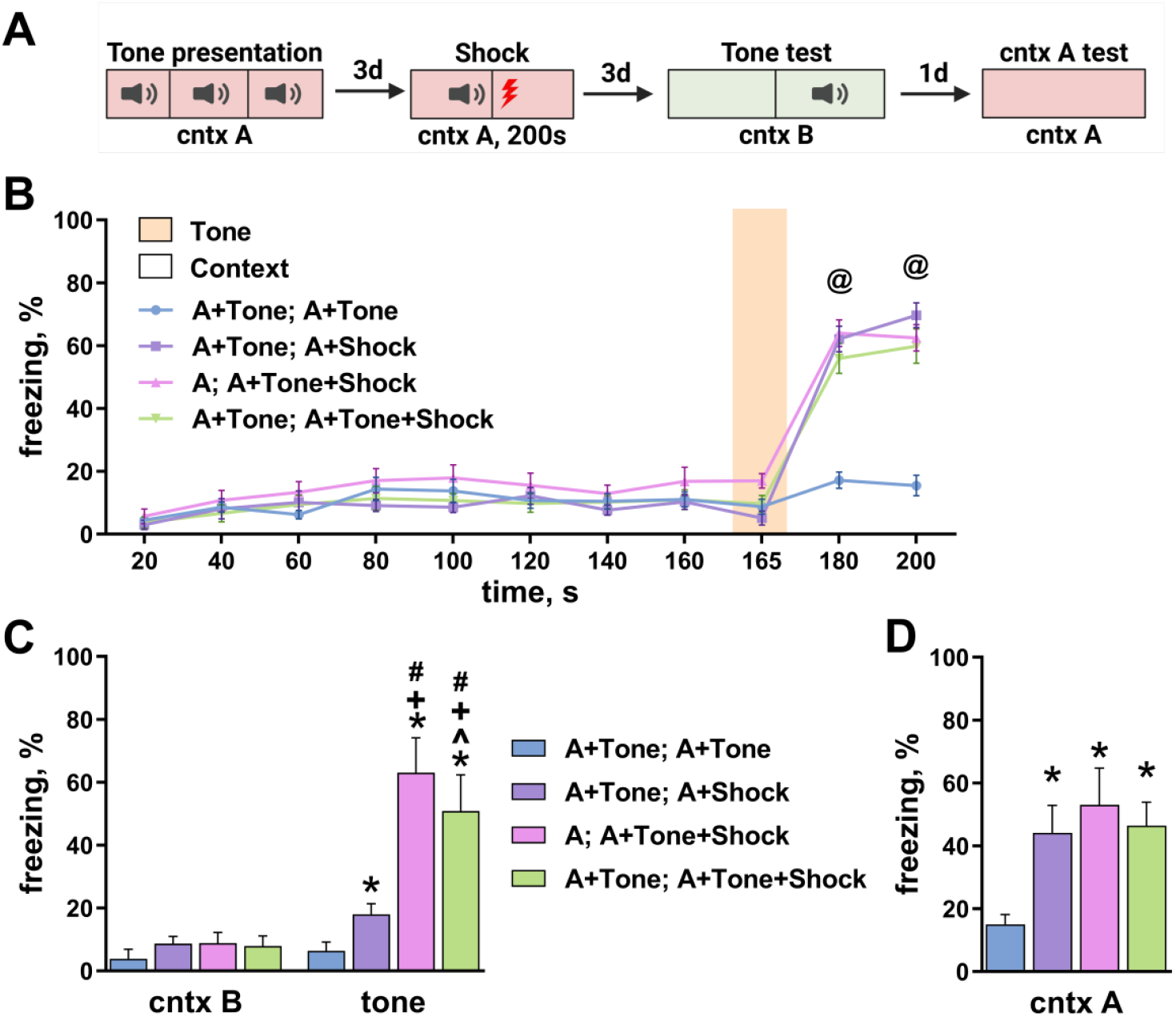
Longer context exploration during conditioning enables tone fear and reveals latent inhibition after tone preexposure. (A) Experimental design. (B) Freezing during conditioning in Context A; @ p < 0.0001 for all groups compared to “A+Tone; A+Tone”, Tukey’s post hoc test. (C) Freezing in Context B during baseline exploration and during the tone presentation (tone test); * - p < 0.0003 compared to “A+Tone; A+Tone”; + - p < 0.0001 compared to “A+Tone; A+Shock”; ^ - p < 0.0001 compared to “A; A+Tone+Shock”; # - p < 0.0001 compared to freezing in Context A’ within the same group, Tukey’s post hoc test. (D) Freezing during the Context A test; * - p < 0.05, Dunnett’s T3 post hoc test.

During the tone test in Context B, all shock groups showed significantly higher freezing during tone presentation than the active control (group: F(3,74) = 34.27, p < 0.0001; test episode: F(1,74) = 224.6, p < 0.0001; interaction: F(3,74) = 48.42, p < 0.0001; pairwise comparisons vs. active control: p < 0.0001; Figure 4C). However, the group that received FS without tone during conditioning (A + Tone; A + Shock) displayed relatively low freezing to the tone (18.03%) which was significantly lower than in groups that received tone–shock pairing (A; A + Tone + Shock and A + Tone; A + Tone + Shock; p < 0.0001).

Notably, both groups that received tone–FS pairing showed high freezing to the tone, but freezing was higher in the group without tone preexposure (A; A + Tone + Shock) than in the preexposed group (A + Tone; A + Tone + Shock; p = 0.0237).

During the context A test, all FS groups displayed elevated freezing compared to the active control (F(3,74) = 18.35, p < 0.0001; pairwise comparisons vs. active control: p < 0.0001), with no differences among the FS groups (p > 0.4145; Figure 4D).

Therefore, providing time for context exploration during conditioning enabled robust tone fear even under otherwise constrained tone–immediate shock training. Under these conditions, tone preexposure was not required and slightly reduced subsequent cue fear, consistent with latent inhibition. In contrast, extensive context exposure did not produce latent inhibition of contextual fear.

## 3. Discussion

Across four behavioral paradigms, we identified boundary conditions under which a previously neutral auditory cue becomes linked to an aversive outcome and how this linkage depends on contextual similarity at retrieval. When the tone was paired with an immediate footshock after a minimal (7 s) interval, the procedure did not produce robust conditioned freezing to either the tone or the context. In contrast, when tone representation was established during preexposure and conditioning occurred in a related context, subsequent testing revealed context-dependent expression of tone-evoked freezing, consistent with the idea that retrieval in a novel context can unmask cue-driven fear under specific training histories.

Why did the brief tone–immediate shock protocol fail to yield reliable learning? Immediate shock procedures are known to produce an immediate shock deficit when animals have insufficient time to encode contextual features that support associative binding and later retrieval [17, 30]. Even with a brief CS, effective association typically requires that either the context or the CS provides a sufficiently stable representation to bridge the aversive experience and to support later pattern completion. Our results suggest that, under used parameters, neither a durable contextual representation nor a stable cue representation was available during this ultrabrief episode, leading to weak or absent conditioned freezing.

A central finding was that cue-related freezing depended on the context in which memory was expressed. When the cue had been preexposed and conditioning occurred in a related context, switching to a novel test context could promote tone-evoked freezing. This pattern is compatible with context gating of retrieval [6, 31, 32] and with the broader framework that memories can be linked while retaining their identity through overlap or coupling of engram ensembles [23, 24, 33]. One possibility is that tone preexposure establishes a weak, non-aversive trace that is preferentially reactivated during conditioning in a related environment, whereas retrieval in a sufficiently distinct context reduces competition from contextual fear and allows cue-driven responses to dominate.

Extending the duration of context exploration before conditioning markedly changed the balance between cue and context learning. Longer exploration is expected to reduce the immediate shock deficit by enabling richer contextual encoding and by facilitating hippocampus-dependent representations that support contextual fear [34, 35]. Under these conditions, the tone paired with immediate shock elicited more reliable cue freezing. However, prior tone preexposure now tended to attenuate cue conditioning, consistent with latent inhibition [12, 13]. Thus, the same preexposure manipulation can either facilitate cue-shock linkage or suppress it, depending on how much contextual information is acquired during conditioning.

Notably, extensive context exposure did not simply add context fear to the behavioral phenotype: it reshaped which memory component was expressed across contexts. This asymmetry likely reflects differences in representational dimensionality and competition at retrieval. Contextual fear relies on high-dimensional, configural hippocampal or cortical representations and their interaction with amygdala and prefrontal circuits [4, 6-8], whereas an auditory CS may be retrieved more elementally and can dominate behavior when contextual fear is weak or when the retrieval context reduces contextual pattern completion.

From an engram and memory-allocation perspective, these behavioral effects motivate a mechanistic hypothesis: tone preexposure may bias subsequent engram allocation during conditioning by transiently increasing excitability and/or priming transcriptional programs in subsets of neurons that later become recruited by the aversive episode [23-26]. Such biased allocation could increase functional coupling between the tone trace and the aversive engram when contextual conditions favor co-reactivation, but could also promote competition and reduced associability (latent inhibition) when the cue representation becomes highly familiar and prediction error is low [13]. These processes are likely coordinated across distributed cortical-hippocampal networks, including frontal/medial prefrontal regions implicated in stimulus integration and long-term fear-memory connectomics [36, 37]. Consistent with this view, recent studies of hippocampal representational dynamics across extinction, relapse, and reconsolidation emphasize how contextual state can reshape retrieval and generalization over time [38-40], within a broader engram framework of allocation and access [41]. Recent work further suggests that constrain hyperexcitability and network-level mechanisms that stabilize ensembles and can regulate memory precision and generalization that may be directly relevant to the context-dependent patterns observed here [28, 42].

These interpretations generate testable predictions. First, activity-dependent tagging approaches (e.g., TRAP-based labeling) can be used to quantify overlap between neuronal ensembles engaged by tone preexposure, conditioning, and later retrieval in different contexts [33, 38, 43]. Second, longitudinal calcium imaging could reveal whether preexposure preferentially recruits pre-configured ensembles that later become allocated to contextual threat memories via excitability-dependent mechanisms [25, 44]. Finally, causal manipulations of candidate nodes (hippocampus, amygdala, and medial prefrontal/anterior cingulate circuits) during conditioning versus retrieval could determine whether context similarity primarily gates reactivation/pattern completion or instead alters competition between cue and context components during expression [45, 46].

Several limitations should be noted. First, we quantified conditioned fear primarily via freezing; complementary measures (e.g., risk assessment, locomotor suppression, or autonomic indices) could clarify whether cue-context interactions generalize across response modalities. Second, our contexts were designed to be “similar” versus “novel” based on standard manipulations; future work could parametrize similarity more formally (e.g., graded cue overlap) and directly test how similarity thresholds shape retrieval and generalization.

Overall, the present experiments demonstrate that the linking of a previously neutral sound memory trace and an aversive episode is not a fixed consequence of pairing, but a context-sensitive outcome shaped by the amount of contextual information available during learning and by the retrieval conditions. These findings support a view in which cue and context memories can be flexibly integrated or segregated - potentially through biased allocation and context-gated reactivation of underlying ensembles - yielding distinct patterns of fear expression across environments [6, 31, 32].

## 4. Conclusions

This study identifies behavioral conditions that determine whether a neutral auditory cue memory can be linked to an aversive reinforcement under temporally constrained training. An ultrabrief tone-immediate shock pairing did not produce reliable cue fear, and tone preexposure facilitated tone fear only when retrieval occurred in a context similar to the preexposure/conditioning context. Providing time for context exploration during conditioning enabled robust tone fear even in a novel test context, but revealed latent inhibition after tone preexposure. Together, these findings highlight contextual processing and CS familiarity as key gates for linking and expressing fear memory traces.

## 5. Materials and Methods

### 5.1. Animals

Male C57BL/6 mice (8–10 weeks old; Stolbovaya breeding facility, Russia) were used. Animals were housed five per standard cage with ad libitum access to food and water under a 12 h:12 h light–dark cycle. All experiments were conducted during the light phase. All procedures complied with national regulations (Order No. 267 of the Ministry of Health of the Russian Federation, 19 June 2003) and were approved by the Commission on Bioethics of Lomonosov Moscow State University (Application № 148-a, approved during the Bioethics Commission meeting № 151-d held on 20.04.2023).

### 5.2. Fear Conditioning Apparatus and Behavioral Quantification

Fear conditioning and memory tests were conducted using the Video Fear Conditioning System (MED Associates Inc., Fairfax, VT, USA) controlled by VideoFreeze software (v2.5.5.0; MED Associates). Behavior was video-recorded and freezing was quantified automatically. Freezing was defined as the complete absence of movement except respiration [29].

### 5.3. Experimental Contexts

Two distinct contexts were used. Context A was a conditioning chamber (25 × 31 × 22 cm) with a stainless-steel rod floor and metal walls. It was illuminated by diffuse white light (mean illuminance 87 lux) and the background noise level was 25 dB. The chamber was wiped with 70% ethanol between animals.

Context B was an L-shaped black Plexiglas enclosure with a plastic floor (20.5 × 23 cm) covered with a layer of corn cob bedding (RehofixMK2000, J. Rettenmaier & Söhne GmbH + Co.). It was illuminated only by near-infrared light required for video recording and invisible to mice. The background noise level was 7 dB. The enclosure was wiped with 3% acetic acid solution between animals.

In Experiment 3, tone memory was tested in a modified version of Context A (Context A′) designed to share key features with Context A while still being distinguishable. Context A′ was an L-shaped black Plexiglas enclosure with a stainless-steel rod floor. It was illuminated by diffuse white light (mean illuminance 75 lux) and the background noise level was 25 dB. Between animals, the enclosure was wiped with a peppermint tincture solution in 50% ethanol.

### 5.4. Experimental Design

Across experiments, the conditioned stimulus (CS) was a 5 s tone (80 dB, 9 kHz). The unconditioned stimulus (US) was a 1 mA footshock (FS) lasting 2 s, delivered through the grid floor. Animals were returned to their home cages immediately after the procedure unless noted otherwise.

#### 5.4.1. Is a 5 s Tone–Immediate Shock Pairing Sufficient for Cue Learning?

In the target group (A + Tone + Shock, n = 20), mice were placed into Context A, received the 5 s tone immediately, and were given a FS immediately after tone offset; the entire procedure lasted 7 s. Two control groups were used: A + Tone (n = 24), which received the tone without shock, and A + Shock (n = 16), which received FS without tone. Three days after training, tone memory was tested in Context B: mice freely explored context for 3 min and then the tone was presented for an additional 3 min. Context A memory was tested for 3 min, 24 h after the tone test.

#### 5.4.2. Does Tone Preexposure Facilitate Subsequent Tone Conditioning in Immediate Shock Paradigm?

The target group (A + Tone; A + Tone + Shock, n = 10) was preexposed to the tone in Context A: mice remained in the chamber for 5 min and received 20 s tone presentations at the beginning of minutes 3, 4, and 5. Three days later, mice received conditioning identical to Experiment 1 (a 5 s tone immediately followed by FS).

Three control groups were included: A + Tone; A + Tone (n = 9; active control, no FS during conditioning), A + Tone; A + Shock (n = 9; tone preexposure, FS without tone during conditioning), and A; A + Tone + Shock (n = 10; context exposure without tone during preexposure, then tone + FS during conditioning).

Tone and context memory tests were performed as in Experiment 1 (tone test in Context B).

#### 5.4.3. Is Expression of Tone Memory after Preexposure Context-Dependent?

Groups and training procedures were the same as in Experiment 2 (n = 10 per group). However, the tone test was conducted in Context A′ instead of Context B to evaluate whether cue memory expression depends on contextual similarity to the training context.

#### 5.4.4. Does Longer Context Exploration during Conditioning Reduce Context Dependence?

Groups were the same as in Experiment 2, with larger sample sizes: A + Tone; A + Tone + Shock (n = 20), A + Tone; A + Tone (n = 18), A + Tone; A + Shock (n = 20), and A; A + Tone + Shock (n = 20).

During conditioning, mice explored Context A for 150 s before the 5 s tone. FS was delivered immediately after tone offset, and mice remained in the chamber for an additional 43 s. Thus, total time in Context A during conditioning was 200 s (vs. 7 s in Experiments 1–3). Tone and context memory tests were performed as in Experiment 1 (tone test in Context B).

### 5.5. Statistical Analysis

Statistical analyses were performed using GraphPad Prism 10.1 (GraphPad Software Inc., San Diego, CA, USA). One-way Brown-Forsythe and Welch ANOVA, followed by Dunnett’s T3 post hoc test as well as two-way ANOVA, followed by Tukey’s post hoc test. The significance threshold was set to p < 0.05. Data are presented as means with 95% confidence intervals.

## Author Contributions

O.I.I. and K.A.T. performed the behavioral experiments, O.I.I. and K.A.T. analyzed and interpreted the data; O.I.I., K.A.T. and K.V.A. wrote the manuscript. O.I.I., K.A.T. and K.V.A. reviewed the manuscript. All authors have read and agreed to the published version of the manuscript.

## Funding

The study was conducted under the state assignment of Lomonosov Moscow State University. Institutional Review Board Statement

All procedures complied with national regulations (Order No. 267 of the Ministry of Health of the Russian Federation, 19 June 2003) and were approved by the Commission on Bioethics of Lomonosov Moscow State University (Application № 148-a, approved during the Bioethics Commission meeting № 151-d held on 20.04.2023).

Informed Consent Statement Not applicable.

## Data Availability Statement

Data are available from the corresponding author upon reasonable request. Technologies.

